# Isoflurane produces antidepressant effects and induces TrkB signaling in rodents

**DOI:** 10.1101/084525

**Authors:** Hanna Antila, Maria Ryazantseva, Dina Popova, Pia Sipilä, Ramon Guirado, Samuel Kohtala, Ipek Yalcin, Jesse Lindholm, Liisa Vesa, Vinicius Sato, Joshua Cordeira, Henri Autio, Mikhail Kislin, Maribel Rios, Sâmia Joca, Plinio Casarotto, Leonard Khiroug, Sari Lauri, Tomi Taira, Eero Castrén, Tomi Rantamäki

## Abstract

A brief burst-suppressing isoflurane anesthesia has been shown to rapidly alleviate symptoms of depression in a subset of patients, but the neurobiological basis of these observations remains obscure. We show that a single isoflurane anesthesia produces antidepressant-like behavioural effects in the learned helplessness paradigm and regulates molecular events implicated in the mechanism of action of rapid-acting antidepressant ketamine: activation of brain-derived neurotrophic factor (BDNF) receptor TrkB, facilitation of mammalian target of rapamycin (mTOR) signaling pathway and inhibition of glycogen synthase kinase 3β (GSK3β). Moreover, isoflurane affected neuronal plasticity by facilitating long-term potentiation in the hippocampus. We also found that isoflurane increased activity of the parvalbumin interneurons, and facilitated GABAergic transmission in wild type mice but not in transgenic mice with reduced TrkB expression in parvalbumin interneurons. Our findings strengthen the role of TrkB signaling in the antidepressant responses and encourage further evaluation of isoflurane as a rapid-acting antidepressant devoid of the psychotomimetic effects and abuse potential of ketamine.

## Introduction

Major depression is a highly disabling psychiatric disorder, the most significant risk factor for suicide and one of the biggest contributors to the disease burden worldwide^1,2^. Clinical effects of the classical antidepressants appear slowly and many patients respond to them poorly if at all^3^. Electroconvulsive therapy (ECT) remains the treatment of choice for treatment-resistant depression but repeated ECT produces cognitive side effects. Thus, there is a huge unmet medical need for more efficacious and rapid-acting treatments for major depression.

Discovery of the remarkable rapid antidepressant effect of the *N*-methyl-*D*-aspartate receptor (NMDA) blocker ketamine in treatment-resistant patients has inspired renewed enthusiasm towards antidepressant drug development^4^. However, the psychotomimetic properties and abuse potential limit the clinical use of ketamine^5^. Emerging evidence suggests that the rapid antidepressant effects of ketamine are mediated by α-amino-3-hydroxy-5-methyl-4-isoxazolepropionic acid (AMPA) receptor dependent fast synaptic transmission and the release of brain-derived neurotrophic factor (BDNF) that further leads to increased signaling of the TrkB-mTOR (mammalian target of rapamycin) pathway, alterations in dendritic spine dynamics, synaptic plasticity and animal behaviour^6–8^. Inhibition of glycogen synthase kinase 3β (GSK3β) is another molecular signaling event recently implicated in the antidepressant mechanisms of ketamine^9^. These effects set forth by ketamine co-associate with altered GABAergic interneuron function^10^.

Brief deep isoflurane anesthesia has been shown to produce rapid antidepressant effects in both double-blind and open-label clinical studies^11–14^. The clinical observations have been, however, partially inconsistent (see^15,16^), which has reduced the interest to thoroughly evaluate isoflurane as an alternative for ECT and ketamine. To this end, we have translated the treatment concept into preclinical settings and evaluated its antidepressant-like and neurobiological effects in rodents. We show that a single isoflurane anesthesia rapidly transactivates TrkB, regulates the mTOR and GSK3β signaling pathways as well as activity and function of GABAergic interneurons, facilitates hippocampal long-term potentiation and produces antidepressant-like behavioural responses in rodents. These findings shed light on the neurobiological basis of clinically observed rapid-acting antidepressant effects of isoflurane and may become instrumental in dissecting shared mechanisms underlying rapid antidepressant responses.

## Results

### Antidepressant-like behavioural effects of isoflurane in rodents

The rat learned helplessness model (LH) has good face, construct and predictive validity regarding depression^17,18^. Whereas long-term treatment with classical antidepressants is required to ameliorate the learned helplessness phenotype^18^, ketamine produces essentially similar antidepressant-like behavioural effect in the model already after a single treatment^19^. Consequently, we employed this model to investigate the potential antidepressant-like effects of isoflurane. Indeed, as shown in Figure 1A, already a single exposure to isoflurane anesthesia produced an antidepressant-like effect (reduced failure to escape from the compartment where the animal receives footshocks) in the LH when the phenotype was assessed at 6 days post-treatment.

Chronic pain is strongly associated with depression^20,21^ and previous studies show that persistent neuropathic pain induces behavioural abnormalities typical for depression and anxiety in mice^22^. Thus, we evaluated the antidepressant-like effects of a single isoflurane anesthesia treatment also in this model. As expected, mice subjected to the sciatic nerve cuffing showed depression-like phenotype (delayed latency to eat food pellet) in the novelty-suppressed feeding (NSF) test at 8-weeks post-surgery (Figure 1B). Importantly, the phenotype was reversed in the nerve-cuffed animals exposed to isoflurane at 12 hours before the testing (Figure 1B). The antidepressant-like effects of isoflurane appeared very rapidly in this test compared to classical antidepressants, which require weeks of repeated administration to show similar actions^23^. Of note, isoflurane did not have antinociceptive effects against the mechanical allodynia produced by the model (Figure 1C).

**Figure 1.**
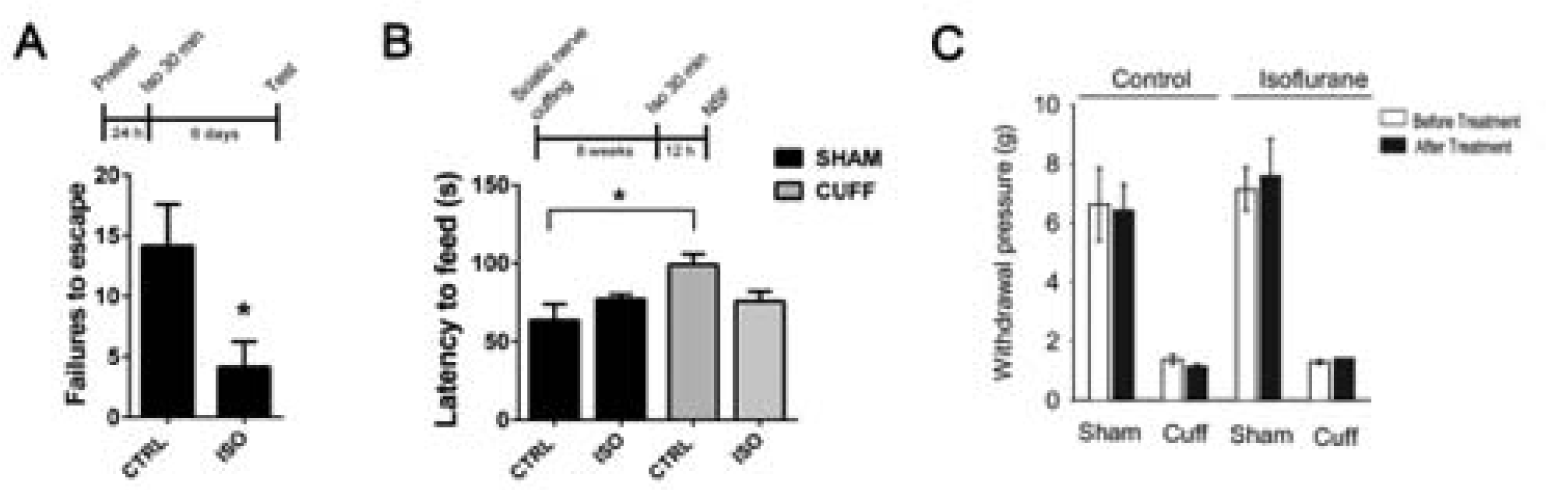
A single brief isoflurane anesthesia produces antidepressant-like behavioural effects in rodents. (**A**) A single isoflurane anesthesia (30 min) decreases the escape failures in the learned helplessness test when tested 6 days after the anesthesia (p=0.0296; n=7). (**B**) Mice subjected to right common sciatic nerve cuffing show anxiodepressive behaviour (increased latency to feed) in the novelty suppressed feeding test. Such phenotype is not seen in mice exposed to a single isoflurane anesthesia (30 min) at 12 hours before testing (two-way ANOVA: cuffing*treatment interaction F_3,27_=6,398, p=0.018; n_sham ctrl_=8, n_sham iso_=8, n_cuff ctrl_=8, n_cuff iso_=7). (**C**) Control and isoflurane-treated mice subjected to the right sciatic nerve cuffing show essentially similar mechanical allodynia in the von Frey test 8-weeks post-surgery (Two-way ANOVA: surgery F_1,24_=123.19, p<0.001, treatment F_1,24_=0.52, p=0.47, surgery x treatment interaction F_1,24_=0.006, p=0.93). The experiment was performed 3 days before and 2 days after control or isoflurane treatment. *p<0.05, **p<0.01; Statistical analysis was done using Student’s t test (A) or two-way ANOVA (B). Abbreviations: CTRL, control treatment; ISO, isoflurane treatment; Sham, sham surgery; Cuff, cuff surgery; NSF, novelty suppressed feeding test.

### Isoflurane anesthesia rapidly activates TrkB receptors

Essentially all antidepressants, including ketamine, have been shown to rapidly, within an hour, activate BDNF receptor TrkB and this effect is proposed to be critically involved in the mechanism of action of antidepressants^7,24–26^. To test the intriguing possibility that isoflurane might bring about similar responses on TrkB, we subjected adult mice to a brief isoflurane treatment and collected brain samples and assayed the TrkB phosphorylation status. Deep burst-suppressing isoflurane anesthesia (induction 4%, maintenance ∼2% *ad* 30 min; see Ref. ^27^), which is shown to elicit rapid antidepressant effects in patients^11,13^, robustly increased the phosphorylation of TrkB (pTrkB) in the medial prefrontal cortex (mPFC) (Figure 2A), hippocampus (HC) (Figure 2B) and the somatosensory cortex (**Supplementary Figure 1**) of adult naïve mice while subanesthetic isoflurane (0.3 %) produced no such effects (Figure 2C). The ability of isoflurane anesthesia to induce pTrkB is extremely fast and transient; a significant increase in pTrkB levels was observed within two minutes from the treatment onset and the TrkB phosphorylation returned close to baseline already at 15 min after the termination of anesthesia (Figure 2D).

When transgenic mice over-expressing flag-tagged TrkB in neurons^28^ were treated with isoflurane, an increase in pTrkB was seen in the flag-antibody precipitated samples (Figure 2E), demonstrating the specificity of the effect on TrkB. Since the transgene expression in this mouse line is directed to postnatal neurons^28^, this result also confirms that the pTrkB response takes place in neurons. Isoflurane treatment increased tyrosine phosphorylation of TrkB at the autocatalytic domain (Y706/7) and at the phospholipase-Cγ1 (PLCγ1) binding site (Y816) but not at the Shc binding site (Y515) (Figure 2E), which is consistent with our previous findings with the classical monoamine-based antidepressants^24,26^. Notably, similar changes in pTrkB were also observed after anesthesia with halothane or sevoflurane (**Supplementary Figure 2**).

**Figure 2.**
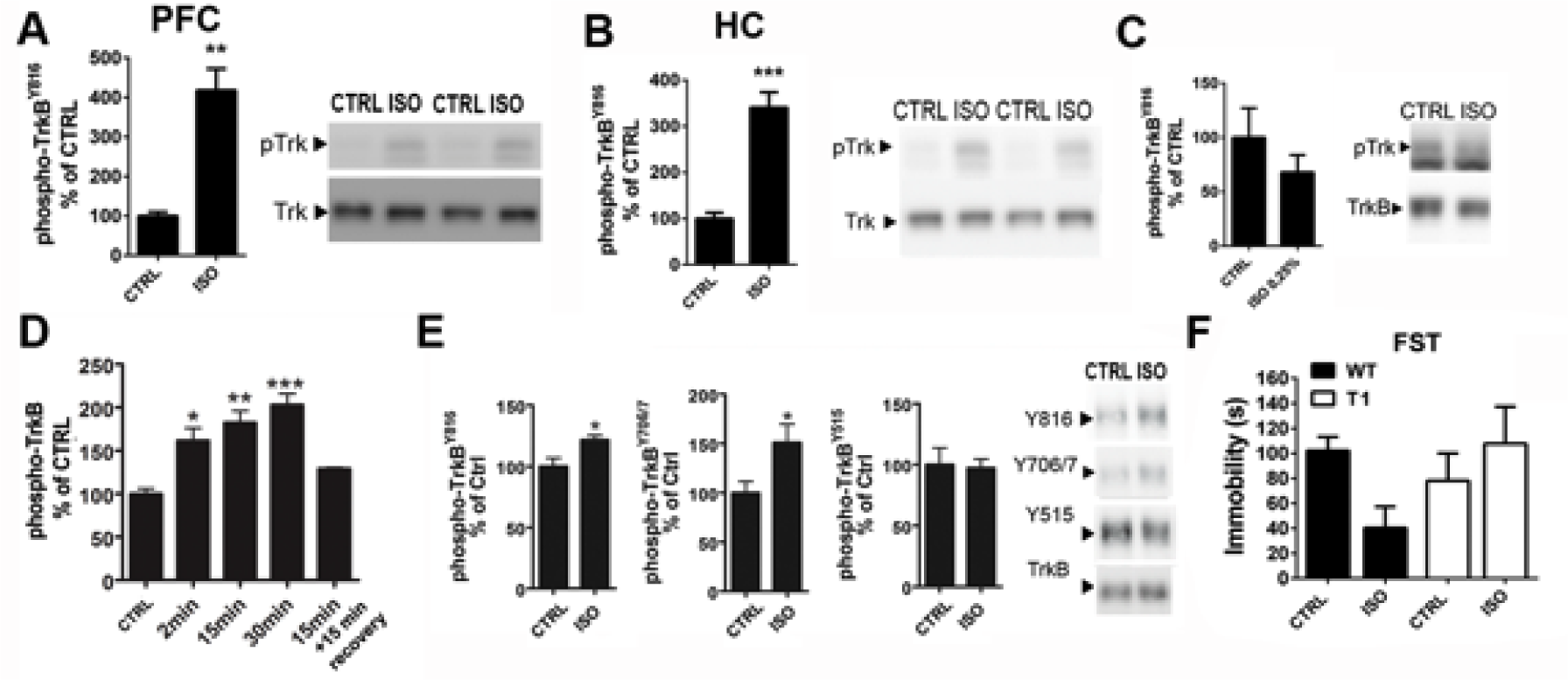
Isoflurane anesthesia induces TrkB autophosphorylation in the brain and produces antidepressant-like effects in the forced swim test. (**A-B**) Isoflurane anesthesia (30 min) increase phosphorylation of TrkB^Y816^ in the adult mouse medial prefrontal cortex (p=0.0022) and hippocampus (p<0.0001). (**C**) Effect of subanesthetic isoflurane (0.3%, 15 min) on phosphorylation of TrkB^Y816^ in the medial prefrontal cortex. (**D**) Time-dependent effect of isoflurane anesthesia on TrkB phosphorylation in the medial prefrontal cortex. (**E**) Significantly increased phosphorylation of the phospholipase-Cγ1 (PLCγ1) binding site (Y816) (p=0.0182) and the catalytic domain (Y706/7) of TrkB (p=0.0426) is detected after flag-immunoprecipitation from hippocampus of mice overexpressing flag-tagged full-length TrkB receptors in postnatal neurons. No change is detected in phosphorylation of Shc binding site (Y515) of TrkB (p=0.8623). (**F**) Wild-type mice treated with isoflurane for 30 min show reduce immobility in the forced swim test when tested 15 minutes after the end of the treatment, whereas in the mice over-expressing the dominant-negative TrkB.T1 isoform the effect was absent (two-way ANOVA genotype*treatment interaction F_3,24_=4,301, p=0.049, n=7). pTrkB levels normalized to total TrkB (A-D). Representative western blots (A, B, C, E) have been cropped from complete immunoblots shown in **Supplementary Information** file. *p<0.05, **p<0.01, ***p<0.001; Mann Whitney U test, Student’s t test, one-way ANOVA followed by Dunnett’s *post hoc* test (C, all groups compared to the Ctrl) or two-way ANOVA followed by Tukey HSD *post hoc* test (E). Abbreviations: CTRL, control treatment; ISO, isoflurane treatment; FST, forced swim test; WT, wild-type; T1, mice overexpressing TrkB.T1; PFC, prefrontal cortex; HC, hippocampus.

### TrkB activation is necessary for the antidepressant-like responses to isoflurane in the forced swim test

The forced swim test (FST) is an acute behavioural test used for screening of compounds with antidepressant potential in naïve animals^29^. Indeed, mice tested 15 minutes after the termination of isoflurane anesthesia showed significantly reduced immobility in the FST compared to sham-treated animals (Figure 2F). The acute behavioural change produced by isoflurane anesthesia was likely not due to an overall increase in locomotor activity, since isoflurane-treated mice showed reduced rather than enhanced activity in the open field test (**Supplementary Figure 3**). Importantly, the effects of isoflurane in the FST were absent in mice over-expressing the dominant-negative truncated TrkB.T1 isoform (Figure 2F). This finding is consistent with the loss of behavioural effects of classical antidepressants in these mice^24^ and suggests that TrkB is required also for the antidepressant-like effects of isoflurane in the FST.

### Isoflurane regulates intracellular signaling implicated in antidepressant responses

Classical antidepressant drugs have been shown to activate the transcription factor cAMP response element binding protein (CREB) downstream of TrkB^24,26^. Along with this change, ketamine has been shown to activate protein kinase B (Akt) and mTOR signaling pathways and phosphorylate GSK3β to the inhibitory Ser^9^ residue^6,7,9^. Ketamine-induced activation of mTOR further leads to phosphorylation and activation of p70S6k and eukaryotic initiation factor 4E binding protein 1 (4E-BP1). Isoflurane anesthesia produced remarkably similar phosphorylation changes on all these intracellular signaling molecules: the phosphorylation levels of CREB^Ser133^, Akt^T308^, mTOR^S2481^, p70S6K^T421/S424^, 4EBP1^T37/46^ and GSK3β^S9^ were significantly increased at 30 minutes after the onset of isoflurane anesthesia in the mouse mPFC (Figure 3A). Significant phosphorylations of CREB^Ser133^, p70S6K^T421/S424^, 4EBP1^T37/46^ and GSK3β^S9^ were also observed in the HC after 30 min isoflurane treatment, whereas phosphorylation levels of Akt^T308^ and mTOR^S2481^ were not regulated (Figure 3B). Similarly to the pTrkB response (Figure 2D), these isoflurane-induced molecular changes were observed within minutes from the treatment onset (Figure 3C). Collectively these data demonstrate that isoflurane anesthesia very rapidly regulates molecular events intimately associated with rapid antidepressant responses in brain areas implicated in the pathophysiology of major depression.

**Figure 3.**
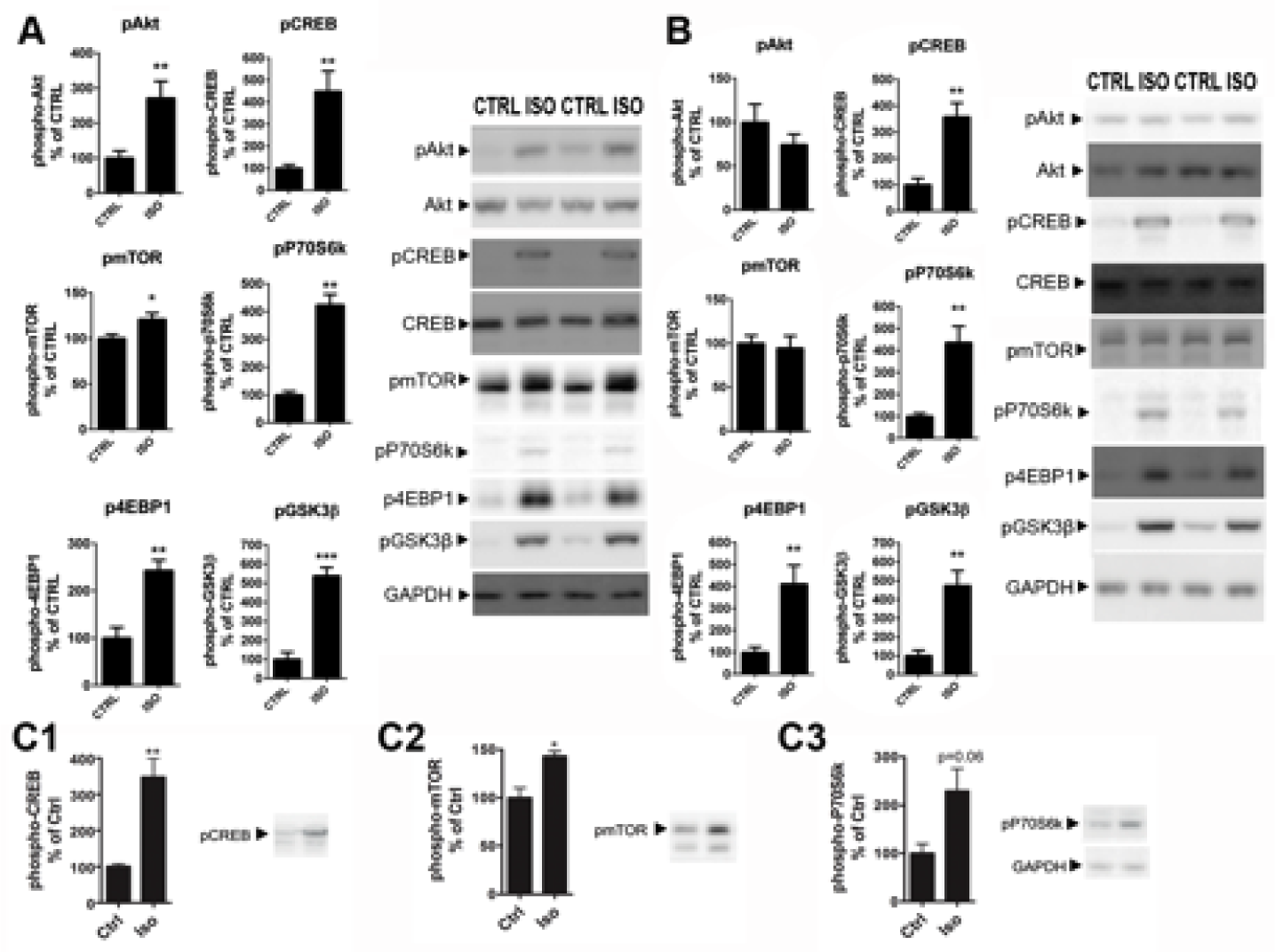
Isoflurane anesthesia regulates intracellular signaling implicated in rapid antidepressant actions. (**A**) Phosphorylation of Akt^T308^ (p=0.0084), CREB^S133^ (p=0.0087), mTOR^S2481^ (p=0.0292), p70S6K^T421/S424^ (p=0.0022), 4-EBP1^T37/46^ (p=0.0012) and GSK3β^S9^ (p=0.0001) in the adult mouse prefrontal cortex after isoflurane anesthesia (30 min). (**B**) Phosphorylation of Akt^T308^ (p=0.3052), CREB^S133^ (p=0.0014), mTOR^S2481^ (p=0.7422), p70S6K^T421/S424^ (p=0.0022) 4E-BP1^T37/46^ (p=0.0022) and GSK3β^S9^ (p=0.0001) in the adult mouse hippocampus after isoflurane anesthesia (30 min). (**C**) Phosphorylation of CREB^S133^, mTOR^S2481^ and p70S6K^T421/S424^ are increased within 2 minutes from the onset of isoflurane (4%) administration. pAKT and pCREB levels normalized to corresponding total protein, other phosphoproteins normalized to GAPDH. n=6/group. Representative western blots have been cropped from complete immunoblots shown in **Supplementary Information** file. *p<0.05, **p<0.01, ***p<0.001; Student’s t test. Abbreviations: CTRL, control treatment; ISO, isoflurane treatment; CREB, cAMP response element binding protein; Akt, protein kinase B; mTOR, mammalian target of rapamycin; GAPDH, glyceraldehyde 3-phosphate dehydrogenase, GSK3β, glycogen synthase kinase 3β.

### Isoflurane transactivates TrkB independently of BDNF and AMPA receptor signaling

Facilitation of AMPA receptor signaling and translation (and release) of BDNF through inhibition of eukaryotic elongation factor 2 (eEF2) are suggested to govern the effects of ketamine on TrkB^6,7,19,30^, yet we found no clear evidence supporting these mechanisms for isoflurane. First, pretreatment with the AMPA receptor blocker NBQX (10 mg/kg, i.p.) for 10 minutes prior to isoflurane had no effect on the isoflurane-induced TrkB phosphorylation and downstream signaling (Figure 4A). Moreover, phosphorylation of eEF2 and levels of BDNF (both mRNA and protein) remained unaltered by isoflurane at the time of TrkB phosphorylation (Figure 4B-D). To investigate the role of BDNF more directly, we performed isoflurane-induced TrkB phosphorylation analyses in mice homozygous for a conditional deletion of BDNF specifically in CamKII-positive neurons. Importantly, isoflurane readily activates TrkB in the hippocampus of these mice (Figure 4E) even though the BDNF protein levels are undetectable (Figure 4D). Notably, essentially similar BDNF-independent acute TrkB (trans)activation has been observed with classical antidepressant imipramine^31^.

**Figure 4.**
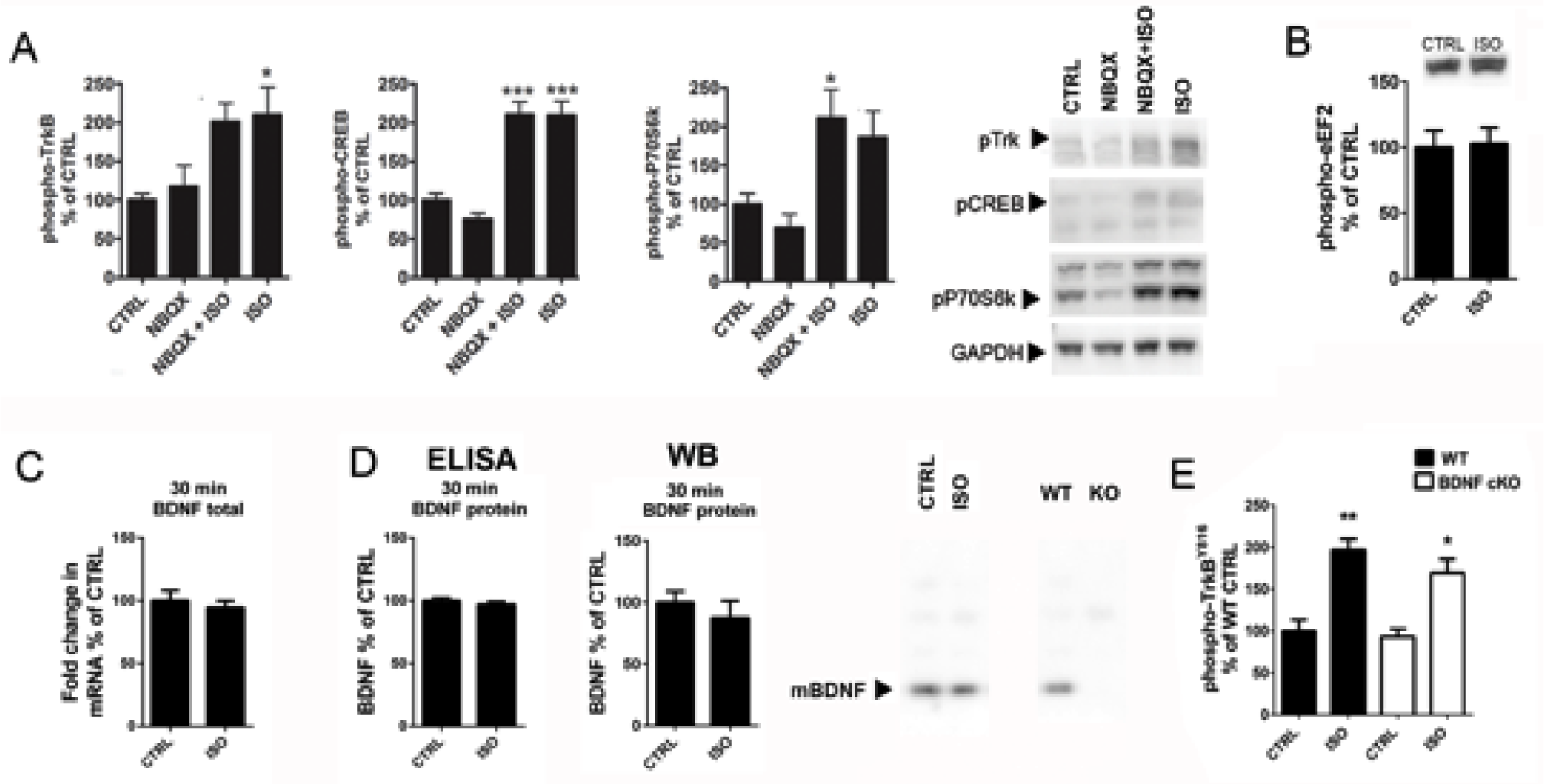
Isoflurane transactivates TrkB. (**A**) Pre-treatment with the AMPA receptor blocker NBQX (10 mg/kg, i.p.) does not prevent isoflurane (15 min) induced phosphorylations of TrkB (two-way ANOVA: F_3,20_=4.379, p=0.017; Tukey HSD *post hoc* test: CTRL vs. NBQX p=0.972, CTRL vs. NBQX+ISO p=0.075, CTRL vs. ISO p=0.050, NBQX+ISO vs. ISO p=0.996), CREB (two-way ANOVA: F_3,20_=24.731, p<0.001; Tukey HSD *post hoc* test: CTRL vs. NBQX p=0.648, CTRL vs. NBQX+ISO p<0.001, CTRL vs. ISO p<0.001, NBQX+ISO vs. ISO p=1.000) or p70S6K (two-way ANOVA: F_3,20_=6.228, p=0.0037; Tukey HSD *post hoc* test: CTRL vs. NBQX p=0.859, CTRL vs. NBQX+ISO p=0.041, CTRL vs. ISO p=0.143, NBQX+ISO vs. ISO p=0.916). pTrkB normalized to total TrkB, other phospho-proteins to GAPDH. n=6/group. (**B**) The phosphorylation of eEF2 was not affected by isoflurane anesthesia (30min) in the prefrontal cortex. n=6. Student’s t test. (**C**) Total *Bdnf* mRNA (normalized to *Gapdh* mRNA) levels remain unaltered after isoflurane anesthesia (30 min) (n_Ctrl_=6; n_Iso_=5). (**D**) Mature BDNF protein levels remain unaltered after isoflurane anesthesia (30 min). Analysis done using ELISA and western blot. A representative western blot on right shows the mature BDNF band, which is absent in a sample obtained from conditional BDNF knockout (KO) mouse. (**E**) Isoflurane anesthesia activates TrkB in the hippocampus of conditional BDNF knockout mice (Two-way ANOVA treatment effect F_3,12_=41,843, p<0.001; Tukey HSD post hoc test WT CTRL vs. WT ISO p =0.001, WT CTRL vs. cKO ISO p = 0.015). Representative western blots (A, B, D) have been cropped from complete immunoblots shown in **Supplementary Information** file. *p<0.05, **p<0.01, ***p<0.001; Abbreviations: CTRL, control treatment; ISO, isoflurane; AMPA, α-amino-3-hydroxy-5-methyl-4-isoxazolepropionic acid; NBQX, 2,3-dioxo-6-nitro-1,2,3,4-tetrahydrobenzo[f]quinoxaline-7-sulfonamide; CREB, cAMP response element binding protein; eEF2, Eukaryotic elongation factor 2.

### Isoflurane does not regulate dendritic spine formation and turnover in the adult mouse cortex

Ketamine has been shown to rapidly – within 24-hours – promote spine recovery in the PFC of stressed rats by enhancing the mTOR-p70S6K signaling pathway^6^. The ability of isoflurane to readily activate this very same molecular pathway prompted us investigating its effects on dendritic spines 24 hours post-anesthesia. However, the spine densities were not significantly different from that found in the control mice in fixed sections of the PFC, HC or the somatosensory cortex (Figure 5). Since fixed tissue preparation could undermine changes in spine dynamics, we used *in vivo* 2-photon time-lapse microscopy to image individual dendritic spines in the same (head-fixed but otherwise freely moving in the Mobile HomeCage^32^) mouse at 24 hours before (control condition), immediately before and at 24 hours after a brief isoflurane anesthesia. Because the PFC is not easily accessible to *in vivo* 2-photon imaging, we focused on the somatosensory cortex, where isoflurane also activates the TrkB-p70S6k signaling pathway (**Supplementary Figure 2**). Spine formation and elimination rates remained unchanged within 24 hours before and 24 hours after isoflurane, indicating that neither spine density nor spine dynamics were significantly affected by isoflurane anesthesia in this region (Figure 6).

**Figure 5.**
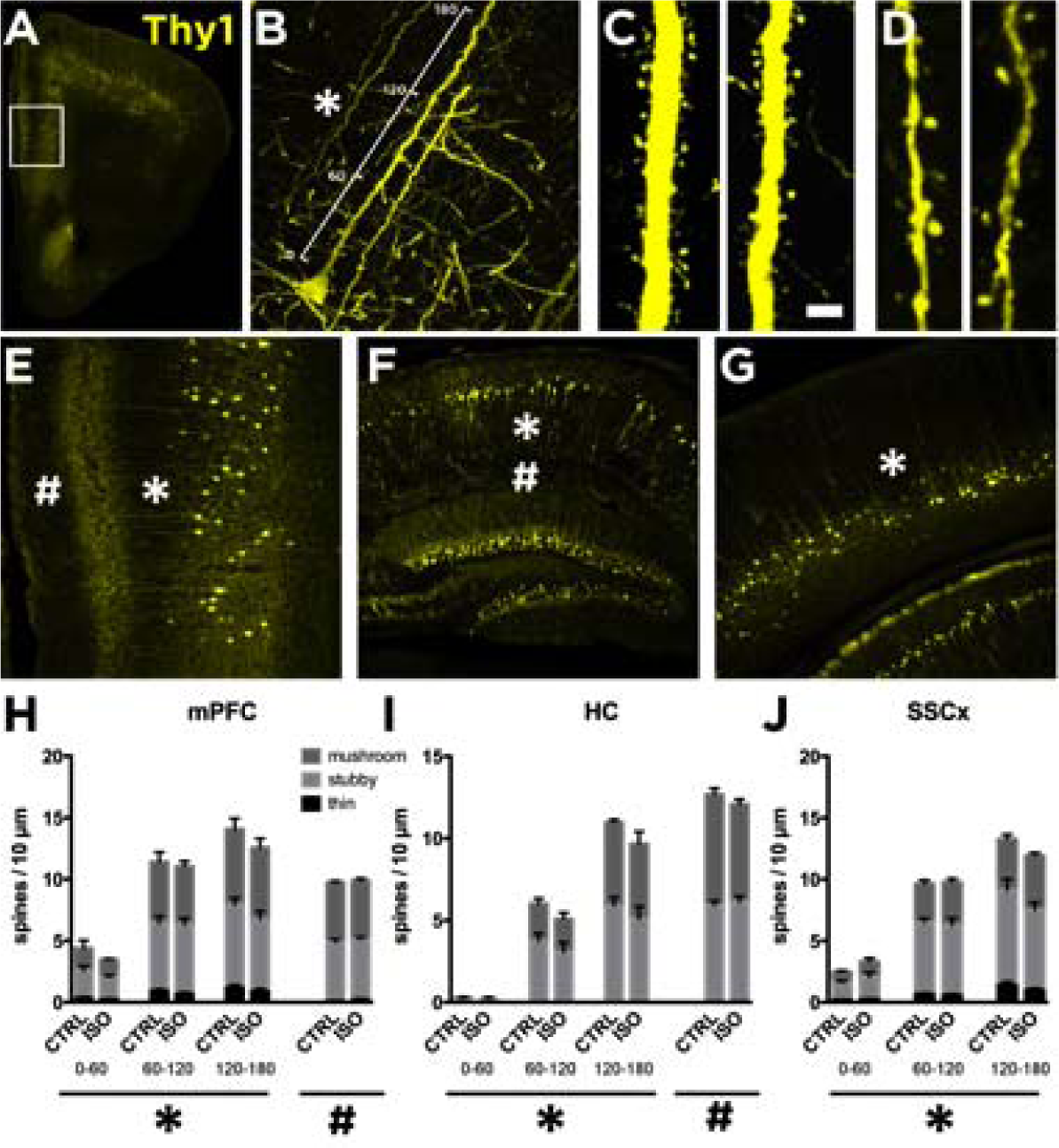
Lack of effects of isoflurane anesthesia on dendritic spines in the mouse cortex and hippocampus. Reconstruction of confocal images showing the medial prefrontal cortex (mPFC) (**A** and **E**), the hippocampus (HC) (**F**) and somatosensory cortex (SSCx) (**G**) from Thy1-eYFP mice. We analyzed dendritic spine density of the primary apical dendrites of pyramidal neurons in three different segments of 60 µm each up to 180 µm from the cell soma (**B**; also indicated with *). Representative images of distal dendritic segment between 120 and 180 µm (**C**). We also analyzed extra-distal dendritic segments in the mPFC and the HC (**D**; indicated as # in **E** and **F**). We analyzed dendritic spine densities from mice subjected to sham or isoflurane treatment (30 min) 24-h before. Histograms showing the densities of spine subtypes in three different segments relative to the distance to the soma (**B**) in the mPFC (**H**; indicated as *), HC (**I**; *) and the SSCx (**J**; *). Isoflurane treatment did not produce changes to the spine morphology or to the total spine amount at different distances from the soma (2-way ANOVA: treatment effect for distance in the mPFC (F_1,36_ =0.7705, p=0.385), the HC (F_1,21_ =1.371, p=0.2547) and the SSCx (F_1,36_ = 0.02270, p=0.8688)). Extra distal segments were also analyzed in the mPFC (**H**; indicated as #; p=0.6756) and the HC (**I**; #; p=0.8203). n=7 mice/group, 3 sections/mouse, 8 dendrites/section. Scale bar in A and D: 4 μm. Statistical analysis was done using Student’s t test (for extra distal segments or # in H and I) and 2-way ANOVA (for the first 180 µm from the cell soma or * in H, I and J).

**Figure 6.**
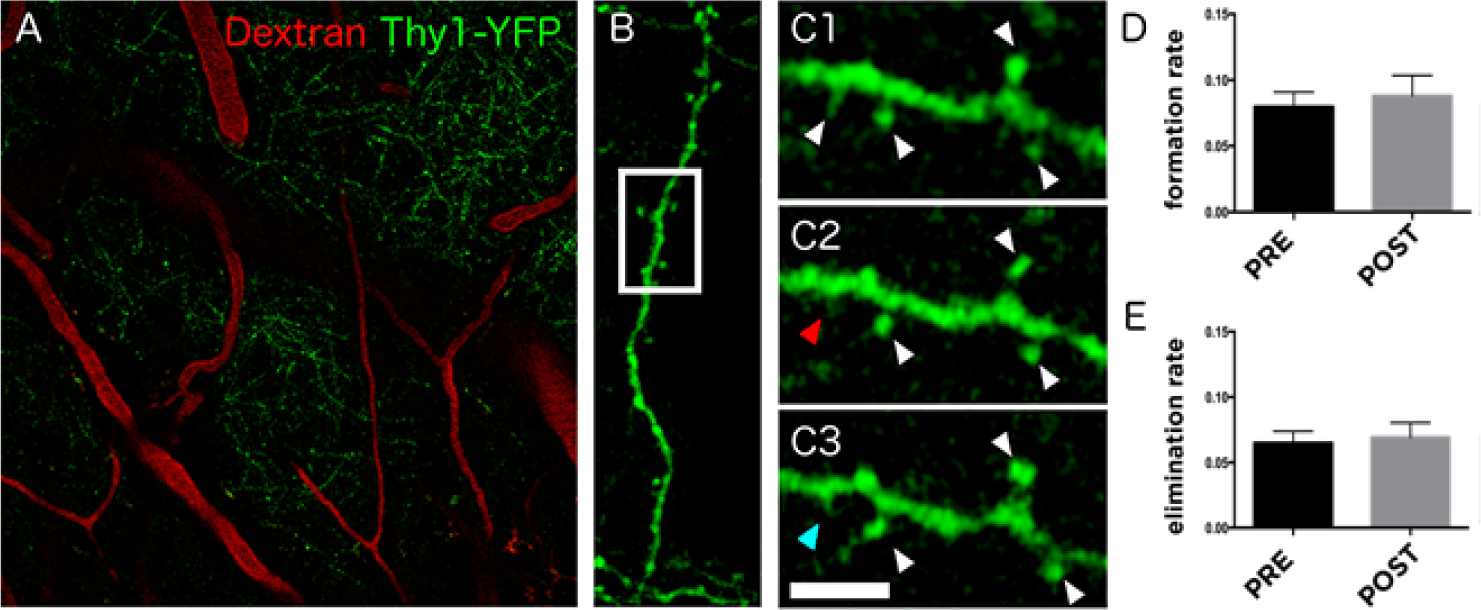
Effects of isoflurane on dendritic spine turnover in the somatosensory cortex of awake mice. (A) Focal plane showing expression of YFP under the *thy1* promoter and the 70 kDa dextran tracer labeled with Texas Red which serves as map to identify the same areas during the different time points of the experiment. (B) Reconstruction of confocal images showing one of the dendritic segments analyzed. (C) Selected area shows a high magnification image of the dendrite to illustrate the spines (C1) 24 hours before, (C2) immediately before and (C3) 24 hours after the isoflurane administration (30 min). White arrowheads indicate stable spines, red arrowhead indicates eliminated spine and blue arrowhead indicates newly formed spine. Histograms represent (D) spine formation (p=0.6476) and (E) elimination rate (p=0.7301) before (pre-treatment) and after (post-treatment) isoflurane treatment. n=3 mice, ∼300 spines/mouse were analyzed. Scale bar size corresponds to 60 μm in A, 11.6 μm in B and 3.7 μm in C images. Statistical

### A brief isoflurane treatment leads to the facilitation of hippocampal synaptic strength

TrkB signaling importantly regulates neuronal plasticity such as long-term potentiation (LTP)^33^. Notably, facilitation of LTP has been implicated in the antidepressant response^5,34^. We therefore used electrophysiology in acute brain slices from mice treated with isoflurane 24 hours earlier to study field excitatory postsynaptic potentials (fEPSP) recorded in the stratum radiatum of the CA1 region of the HC and evoked by electrical stimulation of the Schaffer collateral pathway. Recordings of input–output relationship showed that isoflurane exposure 24 hours before accentuated basal synaptic transmission in the HC (Figure 7A). Paired pulse facilitation was not affected by isoflurane implying unaltered release probability and presynaptic function (Figure 7B). In slices prepared from mice treated with isoflurane 24 hours before, tetanic stimulation (100Hz/1s) produced a significantly higher long-term potentiation (LTP) of the fEPSPs than what was seen in the control slices (Figure 7C) (see Ref. ^35^). These effects of isoflurane on synaptic strength are reminiscent of the previously reported changes seen after a long-term treatment with the classical antidepressant fluoxetine, but they appear much faster^36^.

**Figure 7.**
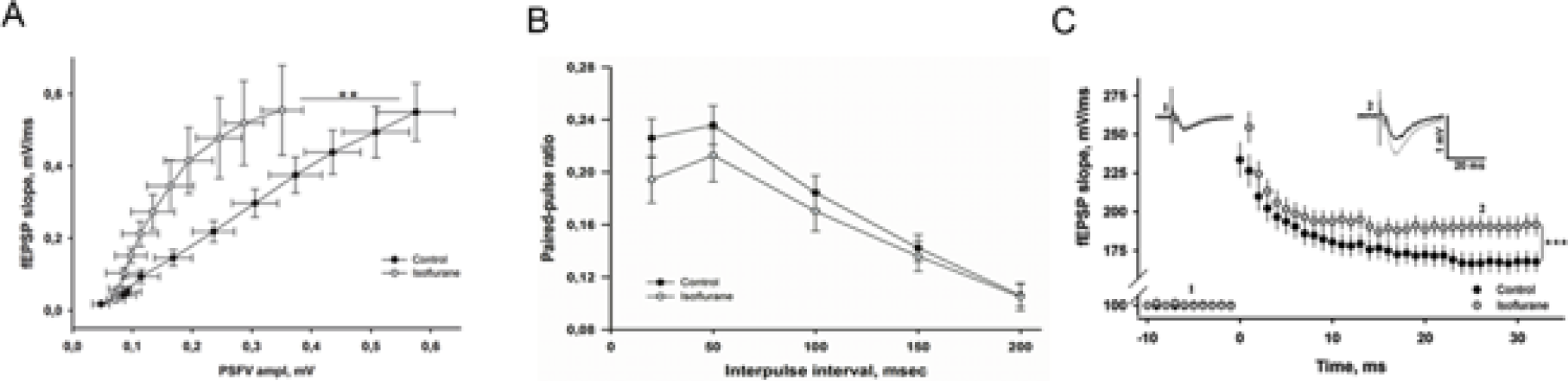
Isoflurane accentuates glutamatergic transmission and plasticity in the hippocampus. (**A**) Slope of field excitatory postsynaptic potential (fEPSP) recorded in the area CA1 plotted as a function of presynaptic fiber volley (PSFV) amplitude recorded in slices from mice treated with isoflurane (30 min) at 24 hours before (n=12 slices/8 mice) and in control animals (n=10 slices/7 mice) (one-way ANOVA: F_1,20_=1.314, p=0.0479). (**B**) Paired-pulse facilitation induced by two consecutive stimuli with interpulse interval (IPI) between 10 and 200 msec was not different between the groups (F_1,10_=0.354, p=0.564). n=9 slices/8 mice for isoflurane group and n=10 slices/7 mice for control group. Statistical analysis was done using one-way ANOVA. (**C**) Long-term potentiation (LTP) induced by high-frequency stimulation (HFS, 100Hz/1s) is significantly enhanced in slices from isoflurane treated animals (two-way ANOVA: F_1,64_=7.91, p=0.000896; n=8 slices/8 mice for isoflurane group and n=8 slices/7 mice for control group). Representative fEPSPs taken before and 30 min after the HFS are shown in the insets (control=black, isoflurane=grey).

### Isoflurane regulates the excitability of parvalbumin-containing interneurons through TrkB

To investigate cellular and network level correlates underlying isoflurane-induced facilitation of synaptic plasticity we performed FosB immunostainings on brain sections with the aim to identify neuronal populations showing sustained activity following isoflurane administration. Interestingly, parvalbumin interneurons within the HC (and cortex; **Supplementary Figure 5**) showed the most prominent FosB immunoreactivity 24 hours after a brief exposure to isoflurane (Figure 8A-B). This observation prompted us to test the role of inhibitory interneurons further using pharmacological and genetic tools. Indeed, bath application of picrotoxin essentially abolished the accentuated LTP after isoflurane anesthesia, supporting a GABA_A_ dependent mechanism (Figure 8C). Moreover, spontaneous inhibitory postsynaptic currents (IPSCs) were significantly increased in CA1 pyramidal neurons in slices obtained from animals treated with isoflurane 24 hours before (Figure 8D). This effect was due to altered excitability of GABAergic interneurons rather than changes in GABA release, since no difference in miniature IPSCs were detected between control and isoflurane treated mice (Figure 8E-F). To investigate the potential involvement of TrkB in these effects produced by isoflurane we performed electrophysiological studies in mice with reduced TrkB expression specifically in the parvalbumin interneurons (PV-TrkB^+/-^ heterozygous cKO mice). Importantly, whereas the effect of isoflurane on spontaneous IPSCs was readily recapitulated in hippocampal slices obtained from control animals, such effect was blunted in slices obtained from PV-TrkB^+/-^ mice (Figure 8G). Altogether these studies suggest that a brief isoflurane anesthesia brings about sustained TrkB dependent changes on the excitability of parvalbumin containing GABAergic interneurons.

**Figure 8.**
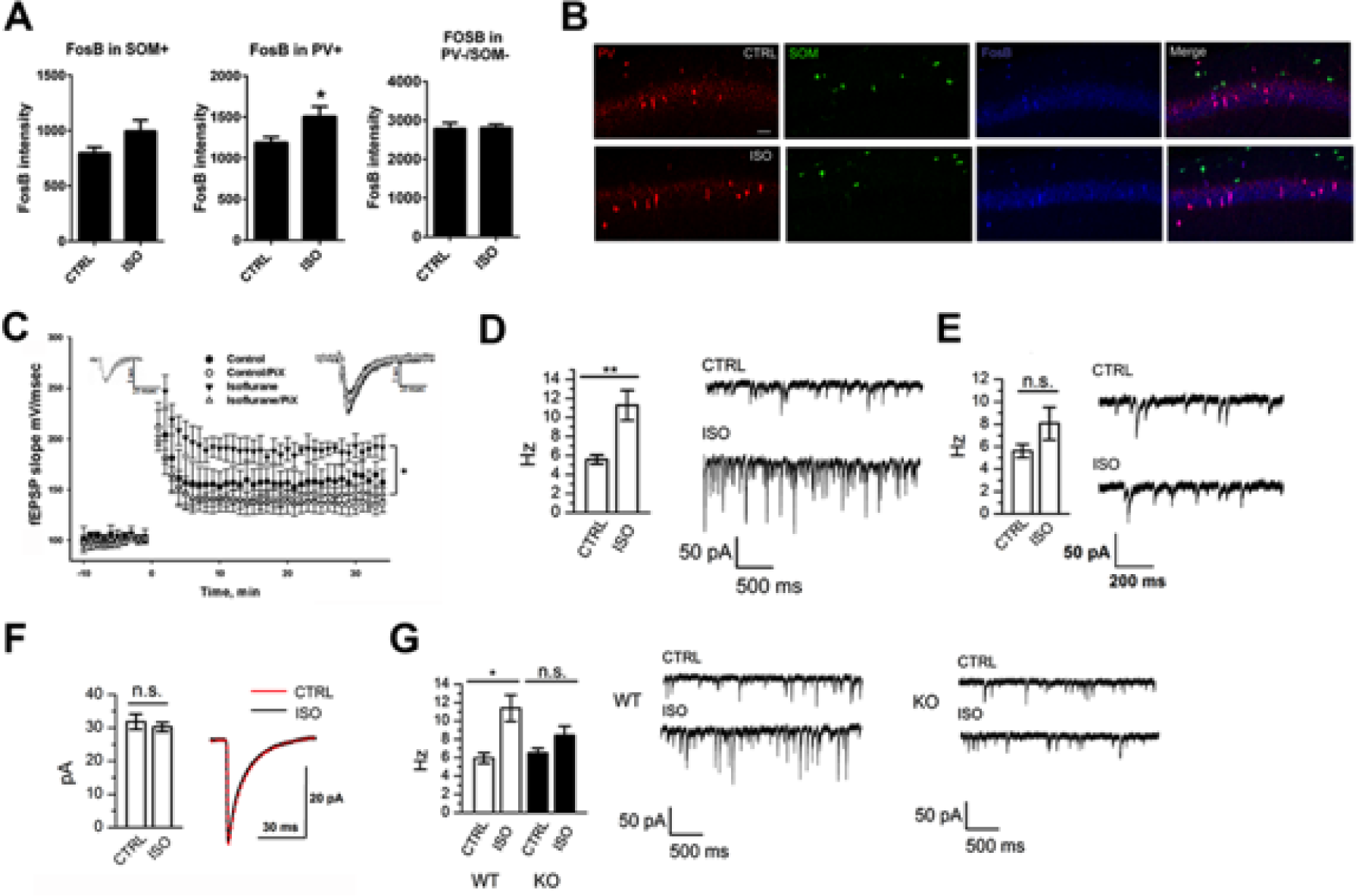
Isoflurane treatment leads to an increase in the activity of parvalbumin positive interneurons *via* activation of TrkB. (**A**) The intensity of FosB staining in parvalbumin positive (PV+) cells, but not somatostatin positive cells (SOM+) or cells not expressing PV or SOM (PV-/SOM-), is significantly increased in the CA1 area of hippocampus of mice treated with isoflurane 24 hours before (p=0.0474, Student’s t test). (**B**) Representative figures of the parvalbumin, somatostatin and FosB stainings. (**C**) Long-term potentiation (LTP) induced by high-frequency stimulation (HFS, 100 Hz) is enhanced in slices from mice treated with isoflurane for 30 min 24 hours before. The difference between the groups disappears in the presence of picrotoxin (PiX). Representative fEPSPs taken 5 min before and 30 min after the HFS are shown in the insets (control=black, isoflurane=dark grey; control/PiX=light gray; isoflurane/PiX=gray). (**D**) The average frequency of spontaneous IPSCs in CA1 hippocampal neurons recorded at 24 hours after isoflurane anesthesia (30 min) (6 slices/6 animals per group, ANOVA, p<0.01) and example traces of the recordings. (**E)** The average frequency of miniature IPSC in CA1 hippocampal neurons (6 slices/6 animals per group, oneway ANOVA) is not affected by isoflurane treatment. (**F**) Decay time distribution of mIPSC (6 slices per group, Pearson’s χ^2^ test). (**G**) The average frequency and example traces of spontaneous IPSC in CA1 hippocampal neurons recorded 24 h after isoflurane anaesthesia in wild-type (WT) and PV-TrkB^+/-^ heterozygous (KO) mice (WT; 6 slices/6 animals for ISO, 5 slices/5 animals for CTRL; KO - PV-TrkB^+/-^; 7 slices/7 animals for ISO, 5 slices/5 animals for CTRL; ANOVA, p<0.05). Abbreviations: CTRL, control treatment; ISO, isoflurane treatment. *p<0.05, **p<0.01

## Discussion

Our study shows that isoflurane, at dosing regimen shown to bring about antidepressant effects in clinical studies^11,13,14^, induces TrkB phosphorylation and downstream signaling in the brain regions implicated in antidepressant responses, and that this signaling is accompanied by long-lasting, TrkB dependent plasticity-related physiological and behavioural responses. Previous observations have suggested that the activation of TrkB and subsequent intracellular signaling are required for the behavioural effects of clinically used antidepressants, including the rapid-acting antidepressant ketamine^7,24^. The observation that TrkB is also activated by and required for the behavioural effects of isoflurane in the FST strengthens the role for TrkB signaling as the common pathway for antidepressant actions.

While many molecular and cellular mechanisms activated by isoflurane resemble those induced by ketamine, there seem to be important differences in the action of these two drugs. It has been proposed that ketamine facilitates AMPA receptor activation, which subsequently leads to BDNF release and TrkB activation. Our data suggest that isoflurane (and classical antidepressants^31^) induces TrkB activation without the involvement of BDNF and AMPA receptor activation^7^. Notably, several neurotransmitters and modulators have been shown to transactivate receptor tyrosine kinases, such as TrkB, through the regulation of Src family kinases^37–39^. Whether such mechanism underlies the isoflurane-induced TrkB activation remains to be investigated. Moreover, isoflurane significantly potentiated glutamatergic synaptic transmission but did not seem to affect synaptic structure (i.e. the morphology and turnover of dendritic spines), while ketamine has been shown to affect both the structure and function of the excitatory synapses^6,30^. The lack of effects of isoflurane on dendritic spine dynamics is consistent with previous observations showing that anesthetics, such as isoflurane, significantly regulate dendritic spine number and morphology during brain growth spurt, but not in the adult brain^40,41^. Unlike ketamine, however, the effects of isoflurane on dendritic spines have been investigated in animals not subjected to chronic stress. Therefore, the effects of isoflurane and ketamine on molecular signaling events, dendritic spines and neuronal plasticity should be investigated and directly compared in naïve animals and in various animal models of depression (e.g. social defeat stress, chronic unpredictable mild stress). The potential differences in the mechanisms of action between ketamine and isoflurane may become instrumental in the attempts to focus drug discovery efforts towards the pathways activated by both compounds.

Reduced concentration of GABA is implicated in pathophysiology of depression and it has been proposed that many of the currently used antidepressants ultimately act through modulation of GABAergic transmission^42,43^. The observation that isoflurane treatment led to changes in the activity of parvalbumin (PV) positive interneurons is consistent with the idea that the primary effect of isoflurane resides in the GABAergic PV interneurons, which regulate the activity of glutamatergic principal neurons *via* perisomatic inhibition^10^. Importantly, tetanic stimulation can result in interneuron-driven GABAergic excitation, leading to synchronous spiking in hippocampal pyramidal neurons and thereby promoting LTP induction^44,45^. Remarkably, clinical studies conducted decades ago have shown that isoflurane therapy that induces burst suppressing pattern in the EEG (electroencephalogram) produces a robust and rapid antidepressant response in a subset of treatment-resistant depressed patients^11,13^. Although some subsequent trials were unable to replicate these findings^15,16^, a recent positive replication by Weeks et al has renewed the interest towards isoflurane as an antidepressant therapy^14^. The antidepressant response of isoflurane appears comparable in magnitude to that produced by the ECT, but faster in onset; a significant antidepressant effect was observed after a single isoflurane treatment^13^. Our current results provide plausible neurobiological basis for the rapid-acting antidepressant efficacy of isoflurane and encourage its further clinical evaluation in depressed patients. This report has significant implications also outside antidepressant field. Isoflurane is a commonly used anesthetic agent in experimental research. With increased use of electrophysiological, imaging and optogenetic methods requiring surgical operations, mice are often exposed to isoflurane anesthesia prior to behavioural testing. Therefore, the long-lasting physiological and behavioural responses to isoflurane treatment uncovered here should be taken into account when designing and interpreting behavioural assessments in animals previously anesthetized with isoflurane. Given the pronounced effects of anesthetics on several molecular pathways in naïve animals^27^, the use of anesthesia should be taken into account in experimental research when interpreting the data.

## Materials and Methods

Detailed description of Materials and Methods are provided in the **Supplementary Information** file.

### Animals

Adult C57BL/6J, BDNF^2L/2LCk-Cre^ (Ref. ^46^), B6.Cg-Tg(Thy1-YFPH)2Jrs/J, TrkB^flox/-^:PV-Cre^+/-^ (Refs ^49,50^) heterozygous conditional knockout mice and transgenic mice over-expressing flag-tagged TrkB or truncated TrkB.T1^28,47,48^ and their wild-type littermates were used. Adult male Wistar rats were used for learned helplessness test. The learned helplessness studies were conducted in conformity with the Brazilian Council for the Control of Animals under Experiment (CONCEA) and were approved by the local Ethical Committee. The experiments involving neuropathic pain model were performed in accordance with guidelines for animal experimentation of the International Association for the Study of Pain (IASP) and European Communities Council Directive 86/6609/EEC and were approved by the local ethical committee of the University of Strasbourg (CREMEAS, n°AL-04). The experiments involving conditional BDNF mutant mice were performed in accordance with the National Institutes of Health Guide for Care and Use of Laboratory Animals guidelines and were approved by the Institutional Animal Care and Use Committee at Tufts University. Rest of the experiments were carried out according to the guidelines of the Society for Neuroscience and were approved by the County Administrative Board of Southern Finland (License numbers: ESLH-2007-09085/Ym-23, ESAVI/7551/04.10.07/2013).

### In vivo pharmacological treatments

The following drugs were used: isoflurane (inhalation; anesthesia: induction ∼4%, maintenance ∼1-2%; see Ref. ^27^), sevoflurane (inhalation; anesthesia: induction ∼8%, maintenance ∼4%), halothane (as for isoflurane) and NBQX (i.p., 10 mg/kg; in saline).

### Immunoprecipitation and western blot

The brain samples were homogenized in NP lysis buffer^26^, centrifuged (16000*g*, 15 min, +4 °C) and the resulting supernatant used for analyses. Samples were separated with SDS-PAGE and blotted to PVDF membrane. Membranes were incubated with antibodies directed against phosphorylated or non-phosphorylated forms of Trk/TrkB, CREB, Akt, mTOR, p70S6K, 4E-BP1, GSK3β, eEF2, BDNF and GAPDH. Following consecutive washes and horseradish peroxidase conjugated secondary antibody incubations immunoreactive bands were visualized using enhanced chemiluminescence using a standard camera.

### qRT-PCR and ELISA

Total *Bdnf* mRNA levels were measured using quantitative RT-PCR as described^51^. BDNF protein levels were analyzed using a commercial ELISA kit.

### Spine analysis from fixed tissue

B6.Cg-Tg(Thy1-YFPH)2Jrs/J mice were transcardially perfused with 4% PFA at 24h after isoflurane/sham treatment under pentobarbital anesthesia. Floating sections (50 μm) were cut using a vibratome and processed for YFP immunofluorescence histochemistry using standard techniques. Images were obtained using confocal microscope and pyramidal neurons from the mPFC and SSCx were selected with the following criteria: intense fluorescence, soma located in layer V, and primary apical dendrite >200 μm long. We imaged the dendrites in three ∼65 μm long segments (proximal/medial/distal). We distinguished three types of dendritic spines: (i) stubby; (ii) mushroom and; (iii) filopodia/thin.

### In vivo two-photon microscopic imaging in awake mice

B6.Cg-Tg(Thy1-YFPH)2Jrs/J mice and Mobile HomeCage system (Neurotar Ltd., Finland) were used for the analysis of dendritic spine turnover in SSCx (layer 1) in an awake *in vivo* imaging through cranial window implanted onto the skull above the somatosensory cortex. Five different dendritic segments from 4 different areas were analyzed per animal to study the time-lapse spine turnover^52^. The spine formation and elimination rates were calculated as the number of spines that have appeared or disappeared, respectively at a given time point compared to the previous time point relative to the total number of spines present at that or the previous time, respectively.

### Immunohistochemistry

The tissue was processed as free-floating sections. After blocking and permeabilization the sections were incubated for 48h at 4°C with a cocktail containing IgG mouse anti-Somatostatin, IgG guinea pig anti-Parvalbumin and IgG rabbit anti-FosB. After washing the sections were incubated with a cocktail containing the fluorescence conjugated secondary antibodies. Washed sections were mounted on slides and coverslipped using fluorescence mounting medium (Dako).

### Electrophysiological recordings

Hippocampal slices (400 μm) were cut with a vibratome as described^53^. Extracellular recordings from CA1 stratum radiatum were obtained with aCSF (artificial cerebrospinal fluid) filled glass microelectrodes (2–5 MΩ) using an Axopatch 200B amplifier. Field excitatory potentials (fEPSP) were evoked with a bipolar stimulating electrode placed in the Schaffer collateral pathway. LTP was induced by 100-Hz tetanic stimulation for 1 s. The level of LTP was measured as a percentage increase of the fEPSP slope, averaged at a 1-min interval 30 min after the tetanus, and compared to the averaged baseline fEPSP slope. For paired-pulse stimulation, interpulse intervals from 10 to 200 ms were tested. To antagonize fast GABA_A_ synaptic transmission picrotoxin (PiX; 100 µM) was used. WinLTP (0.95b or 0.96, www.winltp.com) was used for data acquisition^54^. To investigate inhibitory postsynaptic currents, 25 µM MK-801, 5 µM L-689-560 and 20 µM NBQX were included in the aCSF to antagonize NMDA and AMPA receptors. Whole cell voltage clamp recordings (-70 mV) from CA1 neurons were obtained with glass microelectrodes (4.5 - 6 MOhm) using a Multiclamp 700A amplifier (Axon Instruments, USA). To record miniature IPSCs, 1 µM TTX was added to aCSF. The data analysis was made with Mini Analysis Program, version 5.6.6 (Synaptosoft, USA).

### Behavioural tests

#### Learned Helplessness test

The test was performed as described^55^. The protocol includes a pre-test session (PT) with 40 unescapable footshocks (0.4 mA, per 10s) and on the 7^th^ day a test session (T) in which 30 escapable footshocks (0.4 mA, per 10s) are preceded (5s) by an alarm tone (60 dB, 670 Hz). In the test session the shock could be interrupted or avoided by the animal if it crosses to the other compartment of the chamber during the tone presentation or during the footshock application. Twenty-four hours after the PT the animals were anesthetized with isoflurane.

#### Neuropathic pain model of depression

Neuropathic pain was induced by placing a cuff around the main branch of the right sciatic nerve under anesthesia^22,56^. Sham-operated mice underwent the same procedure without cuff. The mechanical threshold of hindpaw withdrawal was evaluated using von Frey filaments^56,57^. To assess anxiodepressive phenotype the novelty suppressed feeding (NSF) test was done at 12th hour after the isoflurane/sham treatment^22,23^.

#### Forced swim test

A mouse was placed into a glass cylinder (diameter 19 cm, height 24 cm) filled with water (21±1°C) to the height of 14 cm. Overall immobility was measured during the 6-minute testing period.

#### Open field test

General locomotor activity was investigated for 30 min in an illuminated (300 lux) transparent cage (length 28.5 × height 8.5 × width 20 cm). Interruptions of infrared photo beams were used to calculate the overall distance traveled (cm).

### Statistical analysis

Results are represented as mean ± SEM (standard error of mean) unless otherwise stated. For statistical analysis two-sided tests including unpaired two-tailed Student’s t-test, Mann Whitney U test (non-normally distributed data), one-way analysis of variance (ANOVA), mixed model ANOVA, two-way ANOVA, Pearson’s χ^2^ test and Spearman’s correlation test were used. Levene’s test was used to define the equality of variances. If the variances differed significantly non-parametric test was used. *Post hoc* analysis was conducted with Tukey HSD or Dunnett’s test. Statistically significant p value was set to ≤ 0.05. Outliers were considered as values differing more than 2x standard deviation from the mean of the group. To calculate the proper sample sizes we used power calculations (α=0.05 and power 0.80) and estimated the standard deviations and effect sizes based on our previous experience and literature.

## Acknowledgments

The authors would like to thank Outi Nikkilä, Nina Karpova, Vootele Võikar, Mikko Airavaara, Jussi Kupari, Yasmin Hohn, Sissi Pastell, Maria Partanen and Virpi Perko for technical assistance. This study has been supported by grants from the European Research Council (iPlasticity, # 322742) (E.C.), Sigrid Jusélius Foundation (E.C., T.T.), Academy of Finland (E.C., T.R., T.T.), Orion-Farmos Research Foundation (H.An., S.K.) and Doctoral Program Brain & Mind (H.An.).

## Author contributions

H.An., I.Y., E.C. and T.R. designed the experiments, H.An., P.S., M.R., D.P., I.Y., P.C., S.K., R.G., J.L., H.Au., L.V., M.K., V.S., S.J., L.K., T.T., T.R., J.C., S.L., performed the experiments, H.An., D.P., R.G., T.T. performed statistical analyses, H.An., E.C. and T.R. wrote the paper.

## Conflict of interests

L.K. is a paid employee in Neurotar Ltd.

